# Phylogenetic diversity and regionalization of root nodule symbiosis

**DOI:** 10.1101/2023.09.08.556918

**Authors:** R.A Folk, M.W. Belitz, C.M. Siniscalchi, H.R. Kates, D.E. Soltis, P.S. Soltis, R.P. Guralnick, L.M. Borges

## Abstract

**Aim:** Here we determine centers of species richness (SR), relative phylogenetic diversity (RPD) and centers of paleo- and neo-endemism, and regionalizations of phylogenetic diversity in the mimosoid clade of the legumes to understand the distribution and environmental associates of mimosoids lacking RNS (root nodule symbiosis).

**Location:** Global.

**Time period:** Present.

**Major taxa studied:** Mimosoid legumes.

**Methods:** We built a phylogenetic tree of 1313 species and high-quality species distribution models for 1128 species representing the phylogenetic breadth of the mimosoid clade to identify the geographic distribution of RNS. Centers of significant RPD and endemism were identified using a randomization approach, the latter using CANAPE. Phylogenetic regionalization used a distance-based phylogenetic beta-diversity approach.

**Results:** We recognized nine areas of contiguous high SR as distinct SR hotspots. Non-RNS species occur mainly outside hotspots but are closely correlated with high RPD. Absence of RNS was best predicted by high precipitation, and represents multiple independent phylogenetic assemblages in different biogeographic areas.

**Main conclusions:** SR hotspots are partly incongruent with centers of RPD and phylogenetic endemism. Lineages lacking RNS are distributed in SR hotspots in Africa and the Americas, belong to biogeographically separate species assemblages, and are, in most cases, associated with relatively moist tropical environments with low temperature seasonality and high available soil nitrogen.

## INTRODUCTION

Precipitation is one of the main drivers of plant distributions, especially in the global tropics (Gentry, 1995; Pennington et al., 2009; Ringelberg et al., 2023), where dry areas are potential ancient centers of origin for lineages bearing traits that eventually promoted the formation of temperate deciduous forests, fire-prone savannah and scrub, and other harsh habitats characteristic of today’s more seasonal globe (reviewed in Folk et al., 2020). The legumes (Fabaceae), the most speciose dry tropical forest family by a margin of nearly two-fold (Gentry, 1995), are an archetypal group for the study of the link between precipitation and biogeography. Root nodule symbiosis (RNS), a central adaptation in legumes, is intimately related to their predilection for arid open habitats (Sprent, 2007, 2009; Sprent et al., 2017). A series of recent studies primarily focused on legumes (Adams et al., 2016; Doby et al., 2022; Pellegrini et al., 2016; Siniscalchi et al., 2022) has identified water stress as the preeminent driver of the geographic distribution of RNS, species richness, and phylogenetic diversity, reflecting the importance of nitrogen leaf economics for confronting drought challenge (Adams et al., 2016). This relationship suggests an important link between edaphic ecology and precipitation, with the former often seen as secondary in tropical forest ecology (Gentry, 1988).

However, as highlighted by Sprent et al. (2017), among the most vexing questions in legume biogeography is why non-nodulating and facultatively nodulating legumes, i.e., those that do not or need not engage in RNS with rhizobia, are preponderant in the tropics, whereas extratropical areas have almost entirely nodulating legumes. Similarly, the pattern of greater prevalence of non-nodulators in the Eastern Hemisphere remains unexplained (Sprent et al., 2017), and there is no obvious functional reason to expect this biogeographic disparity. Because plant-bacterial symbiosis likely has a profound impact on dispersal, which should limit successful establishment (Simonsen et al., 2017), the processes underlying these biogeographic patterns are likely key to understanding not only the tradeoffs that lead to successful RNS, or lack thereof of, under a variety of environmental contexts (van Velzen et al., 2019) but also how differences in dispersal dynamics of nodulators differ from those of non-RNS plants as a consequence of symbiosis (Parshuram et al., 2023; Simonsen et al., 2017).

Mimosoids (formerly Mimosoideae, sensu Lewis et al., 2005) in particular are some of the most prominent legumes of the semi-arid tropics and subtropics globally, concentrated between 36°N and 36°S and dominant in savannah biomes, particularly in Africa and Australia (Ardley & Sprent, 2021; Yahara et al., 2013). Mimosoids have numerous adaptations to water stress, such as deciduous leaves displaying nyctinasty, wind-dispersed seeds, strong herbivore defense mechanisms, and RNS (Lewis et al., 2005; Ringelberg et al., 2022). However, as in the broader Caesalpinioideae (sensu LPWG, 2017), RNS states are unstable compared to other nodulating lineages. While a single gain of nodulation is ancestral to the mimosoids, four of ten losses found across the whole of the nitrogen-fixing clade occur within mimosoids (Kates et al., 2022). Even though ancestral scenarios of nodulation gain/loss have proven controversial (Doyle, 2011; Griesmann et al., 2018; Kates et al., 2022; Li et al., 2015; van Velzen et al., 2018, 2019; Werner et al., 2014), absence of RNS in mimosoid legumes clearly represents losses under all the foregoing scenarios, making them one of the clearest model clades (Sprent et al., 2017) for understanding under which environments RNS is not essential (van Velzen et al., 2019). This concentration of losses, mostly in species-poor lineages (Sprent, 2007), suggests that mimosoids may have a different and less evolutionary stable form of nodulation, as suggested by de Faria et al. (2022; see also: Estrada-de Los Santos et al., 2018; Lemaire et al., 2016; Thoms et al., 2021). Mimosoids resemble other members of the Caesalpinioideae in the apparent instability of their nodulation condition, but while losses are found throughout the clade, mimosoids form a stark contrast in the much greater prevalence of nodulation overall (Ardley & Sprent, 2021).

A comparison of distribution of RNS and non-RNS legumes is relevant to understanding the ecological contrast between seasonally dry and mesic biomes. Sprent et al. (2017) argue that lack of nodulation in legumes is strongly associated with lowland tropical tree species and reflects lower N limitation in old-growth lowland tropics (Hedin et al., 2009; Vitousek & Sanford, 1986), which agrees with the prevalence of facultative RNS in tropical understories (Barron et al., 2011). By contrast, the woody legume components of tropical dry regions, interpreted as representing the ancestral legume ecology (Lewis et al., 2005; but see Bouchenak-Khelladi et al., 2010; Raven & Axelrod, 1974), are overwhelmingly nodulating (Ardley & Sprent, 2021). RNS has accordingly been interpreted as foundational to the prominent early-successional function of the legumes in tropical dry forests (Costa et al., 2021). Loss of RNS, by contrast, has been interpreted as a response to changing environmental conditions that disfavor the substantial required photosynthetic investment in symbiosis (Griesmann et al., 2018). As with facultative nodulation, lack of RNS has been connected to habitats that either lack nitrogen limitation (Barron et al., 2011) or lack the conditions that favor higher nitrogen investment (McKey, 1994). What is missing are approaches that directly test these proposed verbal models. Here, we directly test questions about why a trait seemingly so beneficial as RNS becomes dispensable in certain lineages by assessing the spatial distribution of non-nodulating strategies in the mimosoids, and the interaction of these strategies with the abiotic environment.

Although large-scale spatial modeling and endemism analyses have been published for important legume clades from different areas of the world (Barros-Souza & Borges, 2022; Dale et al., 2020; Mishler et al., 2014), and one recent work has regionalized global mimosoid diversity (Ringelberg et al., 2023), major questions regarding legume biogeography in light of RNS are still unanswered (Parshuram et al., 2023; Simonsen et al., 2017; Sprent, 2009; Sprent et al., 2017). To investigate the biogeographic regionalization and endemism of mimosoid legumes, and especially to assess how these align with RNS strategies and the environmental factors most closely associated with RNS, we here implement a broad-scale view of the geographic distributions of lineages in the mimosoid clade by generating densely sampled, detailed species distribution models and linking these with a phylogenomic estimate for the clade. With extensive randomization-based significance tests and statistical modeling, we aim to: (1) identify areas of high species richness, phylogenetic diversity, and endemism, and (2) identify biogeographic regions and environmental conditions associated with lineages that lost nodulation. We hypothesize that moist, nitrogen-rich environments are likely to concentrate lineages that have lost nitrogen-fixing symbiosis, but that this is contingent on biogeographical context through regional factors driving endemism and local evolutionary radiations.

## METHODS

### Phylogenetic framework

—Data for the phylogenetic tree came from a recent large-scale sequencing initiative (Kates et al., 2022), in which 100 low-copy nuclear phylogenetic markers were sequenced using a target capture approach (Folk et al., 2021) from samples of the nitrogen-fixing clade of rosid angiosperms obtained from herbarium specimens. Because of the narrower focus of the current analysis than that of Kates et al. (2022), the raw sequence data were reanalyzed in the current study. We selected all available sequences belonging to the mimosoid clade (1,471 samples), plus nine outgroups, chosen based on the tree from LPGW (2017): *Ceratonia siliqua*, *Cercis canadensis*, *Copaifera officinalis*, *Dialium guianense*, *Dimorphandra parviflora*, *Erythrophleum ivorense*, *Gleditsia sinensis*, *Haematoxylum brasiletto*, *Tamarindus indica*.

We trimmed low-quality sequences with Trimmomatic (Bolger et al., 2014) and assembled the sequenced regions with HybPiper (Johnson et al., 2016). The hybpiper_stats.py script was used to obtain basic statistics on the assemblies, including the number of markers for which multiple copies were assembled per terminal. As only 16 of 1,467 terminals had more than 10% of possibly paralogous genes using this assembly pipeline, we used HybPiper’s automatic paralog resolution procedure to select the putative orthologs.

Individual gene alignments were cleaned by removing sites that had more than 50% missing data, using the pxclsq function in phyx (Brown et al., 2017). RAxML-NG (Kozlov et al., 2019) was used to obtain gene trees for the 100 loci, using the GTR+G nucleotide substitution model for all alignments. A species tree was obtained using ASTRAL-III (Zhang et al., 2018). Because pseudocoalescence trees do not have terminal branches, branch lengths were optimized in RAxML v.8 (Stamatakis, 2014) using the -f e option. TreePL (Smith & O’Meara, 2012) was used to calculate divergence times, using three calibration points (root: 66 m.y., Caesalpinioid crown: 58 m.y., mimosoid clade crown: 10.248 m.y.) obtained from Koenen et al. (2021) and Simon et al. (2009), and a smoothing rate of 10,000.

The “genes at 50%” stat from HybPiper (that is, the count of genes that cover at least half of the reference) was used as a guide for removing terminals with low-quality assemblies from the final tree. We also trimmed tips with extremely long branches, spurious placements based on previous analyses, and duplicate species, resulting in a final tree with 1313 tips. Species names follow the World Checklist of Vascular Plants (https://powo.science.kew.org/).

### Occurrence records

—Georeferenced occurrence records for accepted mimosoid genera were downloaded from the following biodiversity discovery platforms: the Global Biodiversity Information Facility (GBIF), iDigBio, and the SouthEast Regional Network of Expertise and Collections (SERNEC). In total, this resulted in 1,596,036 raw records. We then harmonized the names of these raw records with our standardized species list (the “NitFix taxonomy,” Folk et al., 2021) to deal with taxonomic changes that may have resulted in the use of synonyms in the GBIF, iDigBio, or SERNEC repositories. Duplicate records (that is, data points such as specimen duplicates or record duplicates across data providers collected at the same location, by the same collector, on the same day) were filtered to retain only single records. Species retaining at least five occurrence records after these filters underwent a coordinate cleaning procedure that relied on the CoordinateCleaner package (Zizka et al., 2019). Records were removed by CoordinateCleaner if they: 1) had identical latitude and longitude coordinates, 2) were within 500 m of political country or province centroids, 3) were within a 0.5 degree radius of GBIF headquarters, 4) were within 100 m of a record in a global database of ∼10,000 biodiversity institutions including zoos, botanical gardens, herbaria, and museums. Occurrence records with a distance to the nearest conspecific record greater than 5 times the interquartile range of the nearest distance of all other conspecific occurrence records were flagged as potential outlier records. Finally, maps of occurrence records with flagged outlier records were plotted for manual review, and manual occurrence record filtering was conducted where any remaining problems were identified.

### Niche modeling approach

—Here we used a semi-automated niche modeling approach implemented in R with an extensive manual curation strategy that has been reported previously (Abbott et al., 2022; Folk et al., 2023). Importantly, several aspects of modeling are customized on a per-species basis, including accessible areas and predictor variable set. Steps are summarized in brief as follows.

First, the model accessible area was defined by an alpha hull buffered by the larger value of either 75 km or the 80th percentile of distances between nearest occurrence record pairs. Maxent (Phillips et al., 2006, 2017), a machine learning algorithm for species distribution modeling, was implemented on this accessible area using the R implementation in *dismo* (Fick & Hijmans, 2017). The predictor set comprised 13 bioclimatic variables chosen from the WorldClim variable set (Fick & Hijmans, 2017) to minimize collinearity, together with three soil variables (Batjes et al., 2017; depth profiles summarized as in Folk et al., 2023), and two topographic variables (Amatulli et al., 2018). The final full predictor set was (BIO1, BIO2, BIO4, BIO5, BIO6, BIO8, BIO9, BIO12, BIO13, BIO14, BIO15, BIO16, BIO17, nitrogen (cg/kg), sand (g/kg), soil organic carbon stock (t/ha), elevation, and ruggedness), aggregated using an equal area Mollweide projection at approximately a 21-km by 21-km resolution.

Models were first built from the full predictor set using default Maxent settings; on the basis of iterative calculation of VIFs (variance inflation factors), predictors were sequentially removed from the models with VIF > 5 until no predictors met the VIF condition, yielding species-specific predictor sets for downstream steps. Then tuning parameters were optimized using combinations of range multipliers and feature classes available in Maxent using the R package ENMeval (Kass et al., 2021) with specifics reported in (Folk et al., 2023) and model choice either minimizing AICc (almost all models) or maximizing AUC (only models with AUC < 0.7). We converted continuous niche model outputs into binary (presence/absence) outputs by adapting a true skill statistic approach (Abbott et al., 2022), which selects for a thresholding value that balances commission and omission errors.

### Species richness, relative phylogenetic diversity, and endemism

—Biodiverse v. 3.1 (Laffan et al., 2010) was used to calculate three primary biodiversity metrics: species richness (SR), relative phylogenetic diversity (RPD), and relative phylogenetic endemism (RPE). RPD and RPE are rooted in null branch structure hypotheses that facilitate randomization (see next section). RPD was defined following Mishler et al. (2014) as the ratio of phylogenetic diversity (PD, measured as the sum of branch lengths connecting the terminal taxa present in each location) on the original phylogenetic tree divided by that for a phylogeny with the same topology but with a branch length transformation to impose equal branch lengths. High RPD is therefore interpretable as reflecting regions with relatively long phylogenetic branches, as compared to the overall structure of phylogenetic branching connecting all species. Low RPD correspondingly indicates communities with short branches in relative terms. RPE similarly was defined as PE (phylogenetic endemism) for the original tree divided by a transformed tree with equal branch lengths (Mishler et al., 2014), with PE representing the sum of branch lengths connecting terminal taxa in each location, where each branch length is divided by the range size of the terminal taxon or clade (see Rosauer et al., 2009). Thus, RPE is interpretable as a range-weighted version of RPD; high RPE regions reflect regions with range-restricted taxa having branches longer than in the length-unstructured comparison tree, and low RPE regions similarly reflect range-restricted taxa with shorter branches. To segregate the species set by nodulation trait, we calculated the proportion of species in each grid cell that engage in RNS. First, species were matched to a recent comprehensive genus-level database of nodulating species (Kates et al., 2022), and the number of nodulating species was then divided by SR to yield the ratio.

### Randomizations and endemism categorization

—Because raw diversity measures are relatively uninformative without explicit null predictions (Mishler et al., 2014), we again used Biodiverse v. 3.1 (Laffan et al., 2010) to calculate a series of spatially structured randomizations. A total of 100 randomizations were calculated by randomizing species to grid cells while holding constant the richness of each cell and range size of each species. RPD randomizations are straightforward; values of RPD are calculated across the randomizations set to generate a null distribution for each grid cell. A two-tailed test calculated from the randomizations was then used to classify cells into those that are 1) within null expectations, 2) significantly higher, or 3) significantly lower than null distributions. Significance values of *p* = 0.05 were used to distinguish significance of both RPD and RPE randomizations.

RPE randomizations were set up according to the Categorical Analysis of Neo- And Paleo-Endemism (CANAPE) approach of Mishler et al. (2014) to distinguish centers of endemism. CANAPE is a simple algorithm using randomizations and RPE ratios, selecting grid cells that are significantly high (one-tailed test) in either the numerator (original tree) or denominator (transformed tree) of RPE and then using a two-tailed test of the RPE ratio to categorize cells as neoendemics (ratio significantly low), paleoendemics (ratio significantly high), mixture of both types (numerator and denominator significant but not the ratio, i.e., the numerator and denominator have opposing effect directions), or no significant endemism. Neoendemic areas have higher than expected concentrations of range-restricted short branches, paleoendemic areas have higher than expected concentrations of range-restricted long branches, and areas of mixed endemism have elements of both.

### Phylogenetic regionalization

—To understand relationships among centers of mimosoid diversity and loss of RNS in mimosoids in terms of shared lineages, we implemented phylogenetic regionalizations. These tools use quantitative measures, generally based on community distance metrics framed in phylogenetic terms, to examine phylogenetic species turnover and generate hypotheses of “phyloregions” that reflect clusters of areas that share a distinct lineage makeup (Daru et al., 2017). Phylogenetic turnover (phylobeta) was calculated by comparing the lengths of branches of the total phylogenetic tree that are shared and unshared among pairs of grid cells, weighted by the fraction of each species’ geographic range found in that location (Laffan et al., 2016). As implemented in Biodiverse v. 3.1, this yields a dendrogram from which breaks were manually selected to yield well-defined contiguous sets of grid cells (phyloregions). This represents a reanalysis of mimosoid phyloregions as defined by Ringelberg et al. (2023) with an important methodological difference: whereas Ringelberg et al. (2023) directly used occurrence records, here we use species distribution models, which have the important benefit of reducing species range under-commission in areas that are poorly sampled (Chauvier et al., 2022).

### Environmental associates of grid cell metrics

—We implemented a model choice framework to assess the best environmental predictors of three responses: RPD, CANAPE endemism categories, and proportion of grid cell species engaging in nodulation. Compared to previous work focused on climatic variables Ringelberg et al. (2023), we have considered topography and especially soil as important predictors for investigating RNS distribution in particular. Environmental data were summarized by eight predictors chosen following previous work in the clade (Doby et al., 2022; Siniscalchi et al., 2022) and focusing on those most relevant to defining seasonally dry tropical forest: aridity (calculated as the UNEP aridity index following Folk et al. [2020] as a measure of potential water stress), mean annual temperature (Bio1), temperature annual range (Bio7, a measure of temperature seasonality), annual precipitation (Bio12), temperature of the driest quarter (Bio17, a measure of precipitation seasonality), elevation, and three soil predictors: nitrogen content (chosen because nodulation status was under study), soil pH, and soil carbon content (the latter two important for soil fertility and water retention and well-known predictors of plant distribution).

Models were fit according to two classes: generalized linear models (GLMs) and linear mixed models (LMMs). LMMs are a natural framework for handling spatial autocorrelation because cell proximity can be included as a random effect and variance partitioned separately from the fixed effects (primary predictors). Within each model class, the model set was defined to consider either the full predictor set or a reduced set of five predictors based on the highest magnitude of normalized coefficients in the full model. LMM models additionally fit cell centroid latitude and longitude as random effects. Finally, a spatial-only LMM model was fit using only latitude and longitude as random effects, in order to include a no-environment model in the model choice set. Therefore, for each response variable the model set comprised a GLM-full, GLM-reduced, LMM-full, LMM-reduced, and LMM-spatial no environment. For the categorical CANAPE response, endemism significance categories were lumped to yield a binary response (significant/nonsignificant) that was modeled as a logit function. To properly fit the proportional nodulation response, a mixed beta regression model was fit using the R package glmmTMB (Brooks et al., 2017); the 1-inflated response was transformed with the formula (y∗(*n*−1)+0.5)/*n* (where *n* is the sample size) per developer recommendations. To avoid overfit in the more complex mixed beta model, random effects aggregated latitude and longitude to the nearest degree. Due to large sample sizes, model choice used AIC. All continuous environmental fixed effects were mean centered and standardized prior to model fitting.

Plotting of gridded data revealed differences among hemispheres (both Eastern/Western and Northern/Southern) in environmental responses, motivating the investigation of biogeographic domains in the models. In addition to the above procedures, for the proportion nodulating response, an additional random effect was added representing the classification in the phyloregionalization discussed above. The rationale for this analysis was to use a mixed modeling framework to investigate the confounding effect of biogeographic processes on environmental responses (see Results); differing fixed effects on the responses would indicate a conditionality on local biogeographic context, which might indicate differing histories in local radiations among other processes.

## RESULTS

### Dataset results

—The constructed phylogeny covered 1,313 species (39.8% of species-level diversity, 84.5% of genus-level diversity; completion statistics hereafter assume 3,300 species in 84 genera per LPWG [2017]). Species distribution models covered 1,128 species (34.2% of species-level diversity, 81.5% of genus-level diversity). Genus-level nodulation determinations, matched to the phylogeny, covered 1,261 species (96.0 % of the phylogeny). Per-genus sampling proportions are summarized in Supplemental Table S1.

### Species richness

—As recovered in this analysis, there are nine recognized primary hotspots of SR (numbered in Fig. 1a), listed here in approximately descending order of spatial extent following the nomenclature of CEPF (Critical Ecosystem Partnership Fund; https://www.conservation.org/priorities/biodiversity-hotspots) but combining contiguous areas for the purpose of discussion; this will be discussed further below in the context of phylogenetic regionalization analysis. There were three distinct regions in South America. First, a Tropical Andean region in northwest South America comprised the northern Andes and adjacent Amazonian highlands of Colombia and Peru, as well as a small portion of the highlands of southern Central America. Second, east of this region was a hotspot corresponding to the Guiana Shield. Third was an area comprising eastern Brazil, including portions of the Atlantic Forest, Caatinga, and Cerrado. Three further discontiguous hotspots are also present in Australia. The fourth corresponds closely to the currently recognized East Australia hotspot, fifth is the Southwest Australia hotspot, and sixth is a small region in the northern tropical savannah of Australia. Seventh is a hotspot across the dry tropical forest of eastern Africa, eighth is the Guinean-Congo rainforest of west Africa, and lastly a hotspot in southeast Asia. Although SR in Africa and Asia was much lower than in the regions from the Americas, these regions display higher richness in relative terms compared to neighboring areas, are disjunct from other regions of high SR, and were standout areas in terms of RPD (below); thus, African and southeast Asian were recognized based on the sum of diversity mapping presented here.

**Fig. 1.**
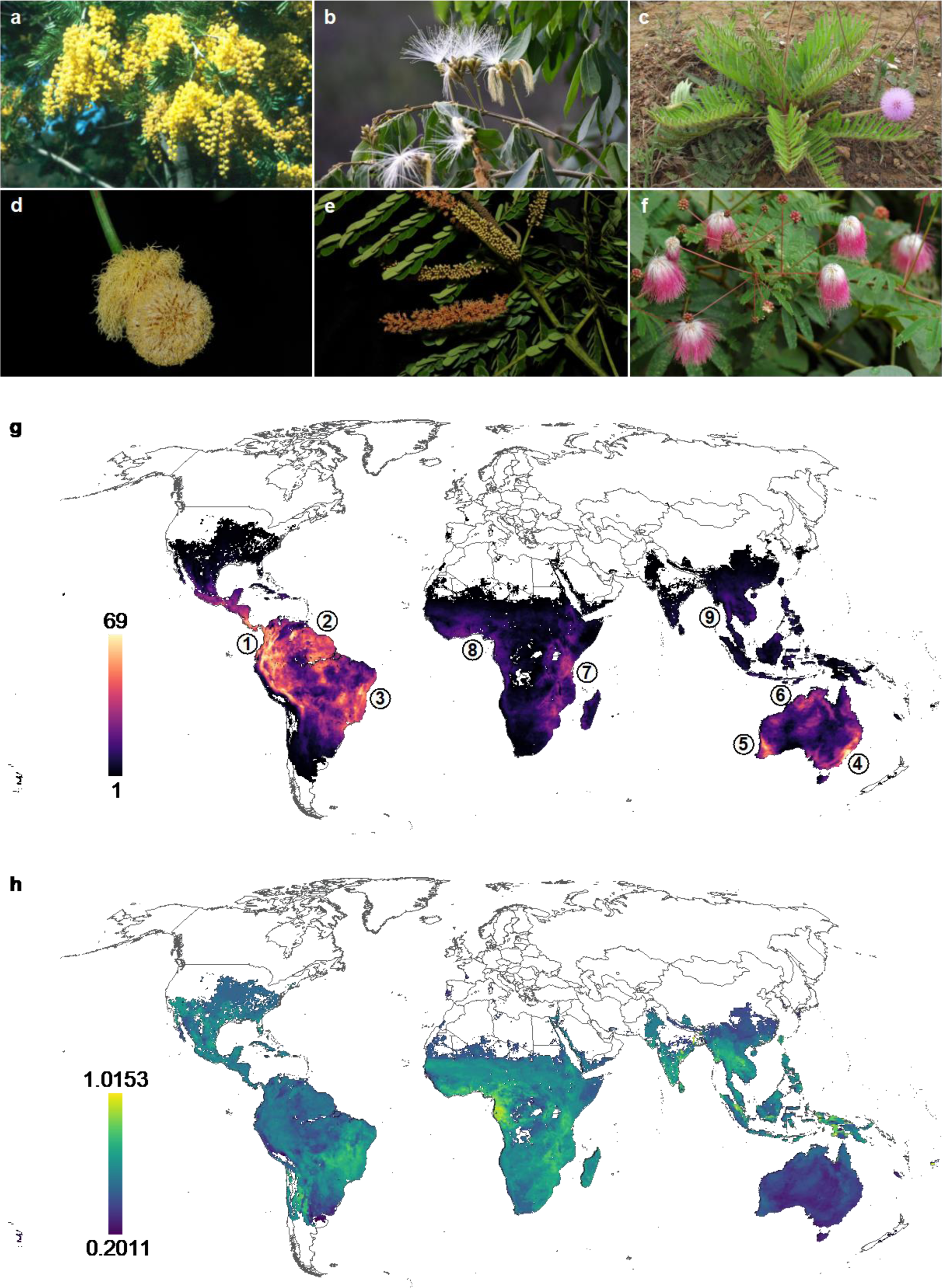
Figure 1. (a)–(c) Examples of large mimosoid genera that nodulate. (a) *Acacia dealbata* Link. (b) *Inga vera* Willd. (c) *Mimosa speciosissima* Taub. (d)–(f) Examples of non-nodulating genera. (d) *Parkia bahiae* H.C.Hopkins. (e) *Tetrapleura tetraptera* (Schumach. & Thonn.) Taub. (f) *Zapoteca caracasana* (Jacq.) H.M.Hern. (g) Mimosoid species richness (SR); warmer red and yellow regions indicate higher SR, with white areas indicating no mapped species. Numbered areas correspond to hotspots discussed in the text; see Results. (h). Mimosoid relative phylogenetic diversity (RPD); warmer yellow areas indicate higher RPD with white indicating insufficient mapped species for the RPD calculation. Photos credits, a,f: Colin E. Hughes; b: Rene Villanueva; c: Leonardo M. Borges; d: Luciano P. Queiroz; e: David Harris.

### Relative phylogenetic diversity

—High RPD regions (Fig. 1b) were similarly delimited with respect to SR, although with different emphases, so the nine regions identified in the previous sections will be followed here. The only major hotspot region in South America for RPD was region three (central-eastern Brazil), which appeared as a contiguous region. The south-central Andes and much of Central America and southern North America also displayed high RPD, but because these areas have low SR, they were not recognized as hotspots above. Similarly, Australia, despite having three clear areas of high SR, had no areas with markedly high RPD.

Africa was the continent with the strongest SR-RPD contrast; both the eastern dry tropical forest (region 7) and the western Guinean-Congo area (region 8) have some of the highest and most spatially extensive mimosoid RPD on Earth, both in absolute terms and compared to null expectations (next section). Thus, these regions are strongly enriched for long branches. Southeast Asia also had markedly high RPD, although this region had the least SR of any of the recognizable hotspots.

### RPD significance areas

—RPD randomizations (Fig. 2a) demonstrated marked spatial structuring of high and low RPD sites. Australia was the only area on Earth with large areas demarcated with low RPD (shorter branches than expected), with significance associated with all three recognized Australia hotspots. All other hotspots were either associated with higher RPD than expected (large areas of Mexico and the southernmost USA, most of Sub-Saharan Africa, southern South Asia, and most of Southeast Asia) or were not significant (northern Andes and the Guiana Shield).

**Fig. 2.**
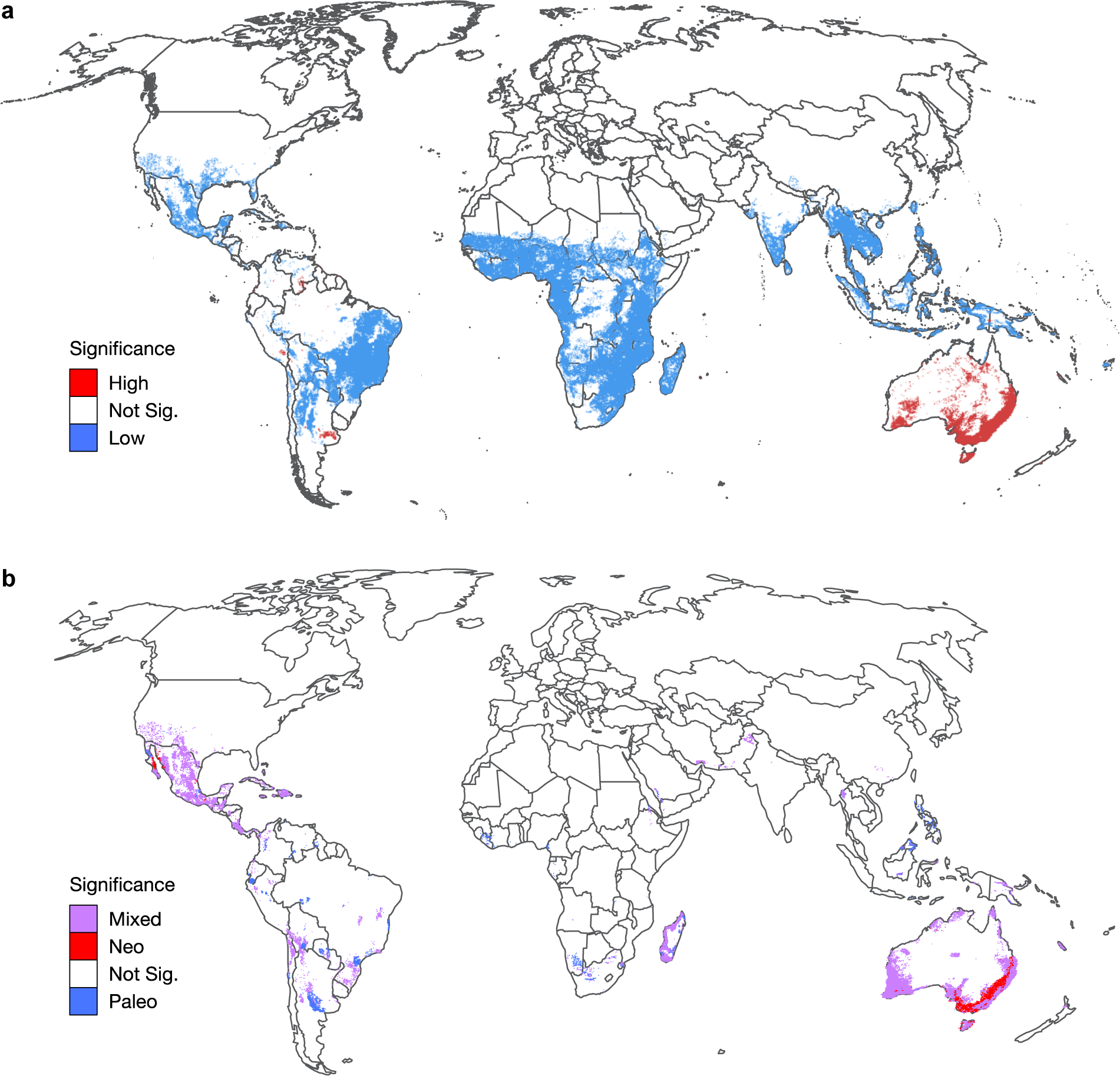
(a) Mimosoid RPD significance. (b) Mimosoid CANAPE significance.

In South America, similar to the results for raw RPD, only the Atlantic Forest-Cerrado region had significantly high RPD; the spatial extent was further westward and included areas of the Chaco and Andean foothills with low SR. Three additional small areas in South America displayed significantly low RPD (red, Fig. 2; see Discussion). Both the eastern and western hotspots recognized in Africa had significantly high RPD, as did the Southeast Asia hotspot.

### Areas of neo- and paleoendemism

—CANAPE analysis (Fig. 2b) demonstrated that only a few of the recognized hotspots of diversity had significant endemism patterns; overall, little signal of endemism was found outside of Australia, Madagascar, and the Americas. In the Americas, the only major significant endemism was in an extensive area including much of the dry interior of Mexico and adjacent southwestern US, identified as mixed endemism. Similar to the RPD analysis, small significant areas approximately corresponding to the tropical-temperate transition were recovered in the middle Andes and Argentina.

In Australia, the Southwest Australia hotspot was identified as an area of mixed endemism, while East Australia was an area of primarily neo-endemism, a typical pattern seen in biogeographic studies of this area. No endemism area was recovered corresponding to the northern region. CANAPE results in Africa and Southeast Asia recovered no large areas of endemism corresponding to the recognized SR hotspots; continental Africa had almost no endemism significance. Coastal western Madagascar, however, was recovered as a center of mixed endemism despite its low SR.

### Phylogenetic regionalization

—The regionalization analysis (Fig. 3) recognized 12 regions that partly followed discontiguous high SR regions that have been recognized above as hotspots. In South America, the Guiana Shield, the Amazon, and the northern Andean highlands were grouped as a region. A further broad region characterized much of the eastern Amazon and included the Atlantic Forest with the Cerrado, Caatinga, and adjacent regions. The Andean region was narrowly defined in the north, but included highland foothills in the southern Andes. A final distinct region comprised temperate grasslands of southern South America. North of this, Mexico and the Caribbean were recognized as a single region, and the southern United States, northern Baja California, and parts of northern Mexico were recognized as the final phyloregion of North America.

**Fig. 3.**
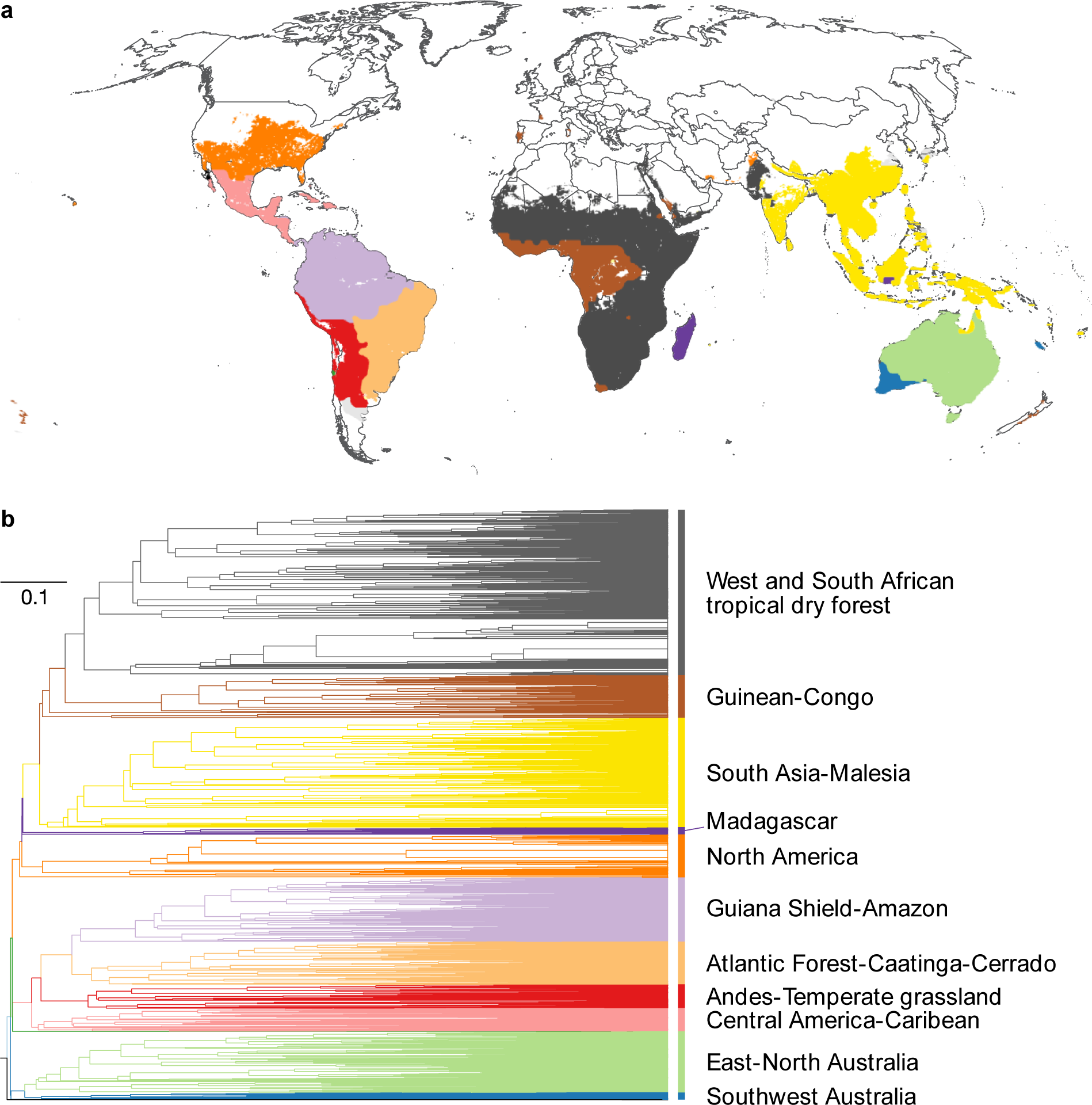
Regionalization analysis. (A) Map of recognized phyloregions. (B) Dendrogram of recognized phyloregions.

In Africa, four primary phyloregions were recognized, with only small areas attributed to adjacent Asian phyloregions. The Guinean-Congo rainforest hotspot was recognized as a region; a small area of southwestern South Africa (the Western Cape) was part of this region. Most of the rest of Africa was grouped with the dry tropical forest hotspot of East Africa as a further phyloregion. Madagascar was clearly delimited as a separate phyloregion.

The three hotspots recognized for Australia were all recovered as separate phyloregions, with the dry interior of Australia attributed to the northern phyloregion. Southwest Australia was recovered as the most distinctive mimosoid phyloregion of all. In Asia, most of Southeast Asia, South Asia, and temperate East Asia were recovered as a single phyloregion that extended south to small areas of rainforest in northern Australia. Of the other regions recovered in Asia, the only other spatially extensive region was the arid zone of northwest India, clustered with the East African phyloregion.

### Distribution of RNS

—Only five biogeographic regions were associated with mimosoid lineages lacking RNS, two of them corresponding to biodiversity hotspots of mimosoids (Fig. 4). First, the richest area on Earth for non-nodulating mimosoids is the Guinean-Congo rainforest hotspot, where grid cells with up to 67% non-RNS species were found. Second, a much smaller proprotion of non-RNS species occurs in the southern part of the dry tropical savannah hotspot of east Africa. Three more regions with non-RNS occurred outside of SR hotspots, all in the Americas: (1) the Amazon, (2) the dry interior of Mexico including the Altiplano and some montane regions, and (3) the northeast of Brazil including areas of Cerrado, in seasonally dry woodlands and other more mesic areas. While RNS had little relationship to SR, non-RNS species were distributed in areas with elevated RPD (compare Fig. 4, Fig.2a).

**Fig. 4.**
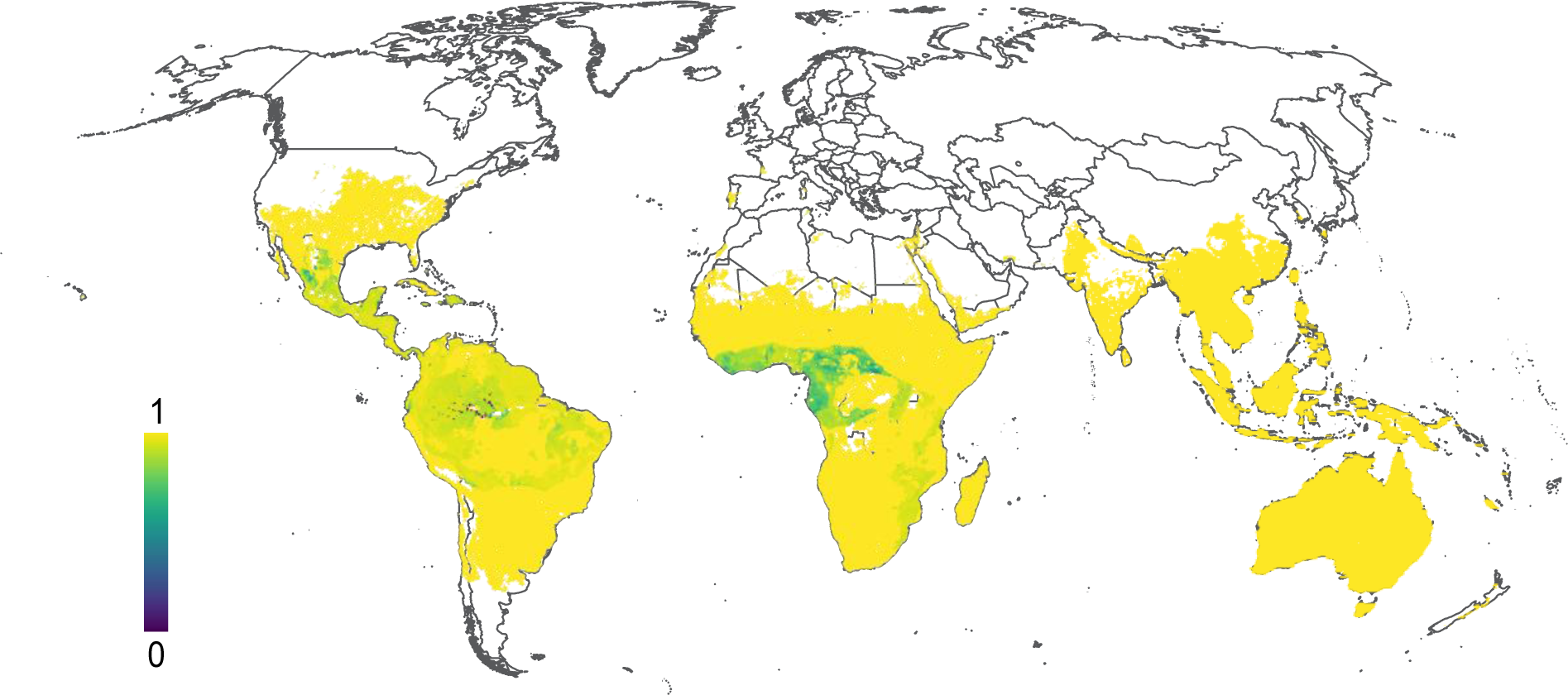
Proportion of nodulating species; cooler blue and green colors indicate a greater number of non-RNS species than RNS species; completely yellow communities indicate the lack of non-RNS species (and thus only RNS species).

### Environmental diversity models

—In the RPD model set, the full (8-predictor) LMM (ΔAIC = 3873.3; Table S2) was strongly favored with moderate predictive power for the fixed effects (marginal *R*^2^ = 0.1517) in a relatively strong model overall (conditional *R*^2^ = 0.7057344). Of the fixed effects, temperature seasonality had the greatest variable importance as measured by standardized coefficients (Table S3), followed by soil pH and aridity. Part of the reason for significant unexplained variation in the RPD model may be that it is confounded by biogeographic province. Investigation of RPD segregated by Hemisphere demonstrated contrasting RPD responses to environment only in the Northern Hemisphere (Fig. 6; not seen in CANAPE significance, below).

In the CANAPE significance model set, the complex LMM was favored (ΔAIC 38.986) and had strong predictive power, with marginal *R*^2^ = 0.0569 and conditional *R*^2^ = 0.9769 (therefore, most of the variance was explained by random effects). This suggested the environmental data had limited power to predict endemism, but a model including fixed effects was strongly favored (ΔAIC = 1014.122 vs. no-environment model). In contrast to RPD, while aridity (Fig. 5a) was in the predictor set, the most important variable was soil pH (endemism was associated with high soil pH), and BIO1 (mean annual temperature) assigned nearly similar importance with significant endemism associated with lower temperature across all significance categories (although the lowest mean, in neo-endemism, ∼15°C, is still subtropical; Fig. 5b). Unexpectedly, higher soil nitrogen (Fig. 5c) was associated with significant endemism, although the data ranged from 25 to 250 cg/kg, and therefore all categories are relatively poor in nitrogen.

**Fig. 5.**
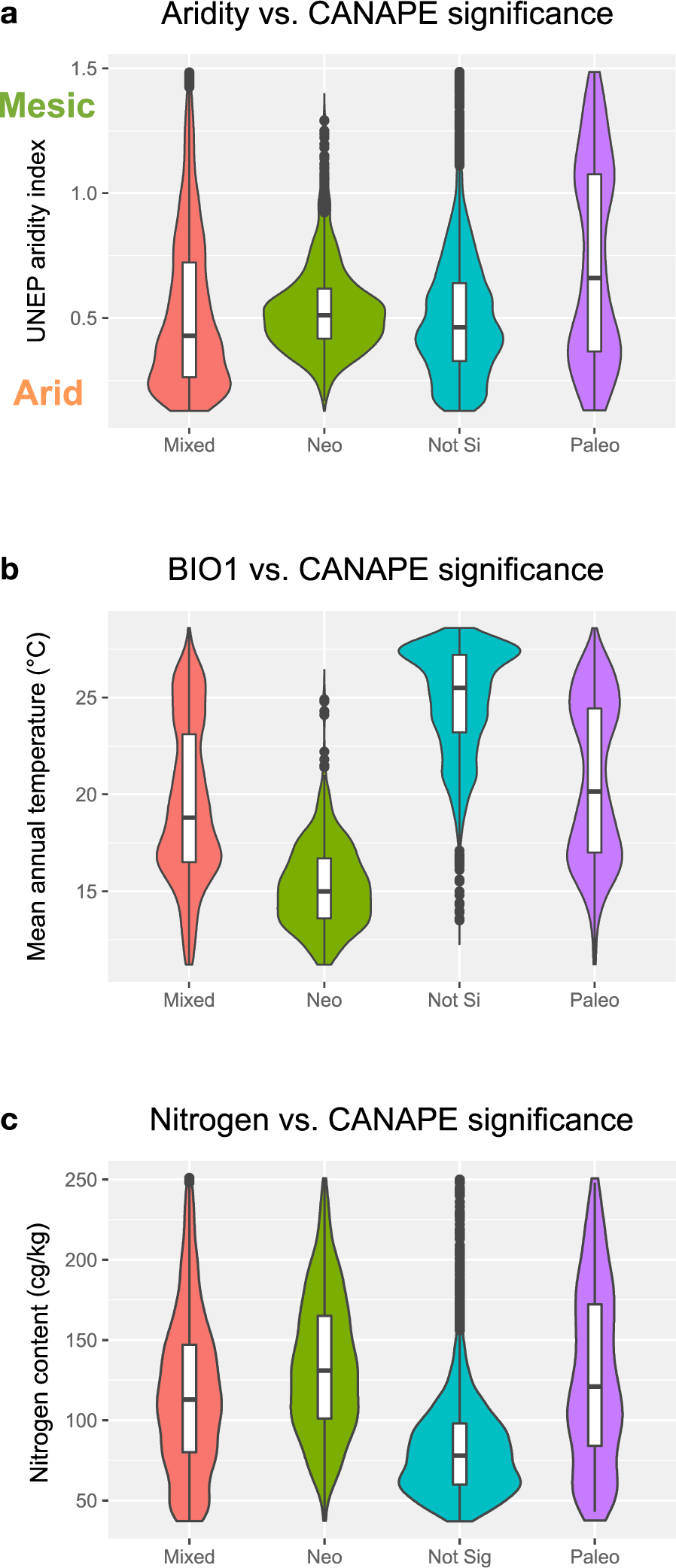
Violin plots of significance categories and non-nodulator presence globally. CANAPE endemism significance categories are shown vs. (A) UNEP aridity index, (B) mean annual temperature, and (C) soil nitrogen content.

The nodulation proportion model instead favored the simple 5-predictor LMM (ΔAIC = 5420.7). The fixed effects had moderate predictive power (marginal *R*^2^ = 0.247). Annual precipitation was the most important variable and aridity was the second most important (similar to results obtained in a continental scale study of all nodulators: Doby et al. [2022] and a study of actinorhizal plants: Folk et al. [2023]), and the direction of the relationship was as expected, with drier environments associated with more nodulating species. Temperature seasonality (BIO7) was also important and was positively associated with nodulating proportion; nitrogen (which was of interest but not in the favored model) was negatively correlated with nodulating proportion as expected, but this is a much weaker effect.

Similar to RPD, mean annual temperature showed a contrasting response between Hemispheres in the nodulating proportion model (Fig. 6), with non-nodulator presence associated with cooler environments in the Northern Hemisphere but warmer environments in the Southern Hemisphere. Thus, the distributions of RNS and RPD, while displaying somewhat differing responses, are both shaped by drought stress and biogeographic regions, reflecting their similar distribution. Because biogeographic domain was of interest, we fit an additional LMM for proportion nodulating that added the phylogenetic regionalization (above) as a third random effect. In this expanded model, the intercepts of the phyloregion random effect can be interpreted as region-specific nodulating effects independent of environment and latitude. The random effect plot (Fig. 7) demonstrates that many regions were close to the mean (zero line), but two regions stand out as outliers below the mean line, suggesting fewer nodulators than expected based on environmental conditions: Mexico (marked “Central America-Caribbean” following Fig. 3) and the Guinean-Congo region. The South Asia-Malesia region was a high outlier, suggesting more nodulators than expected based on the environment. The Central America-Caribbean region was the only significant outlier (Fig. 7).

**Fig. 6.**
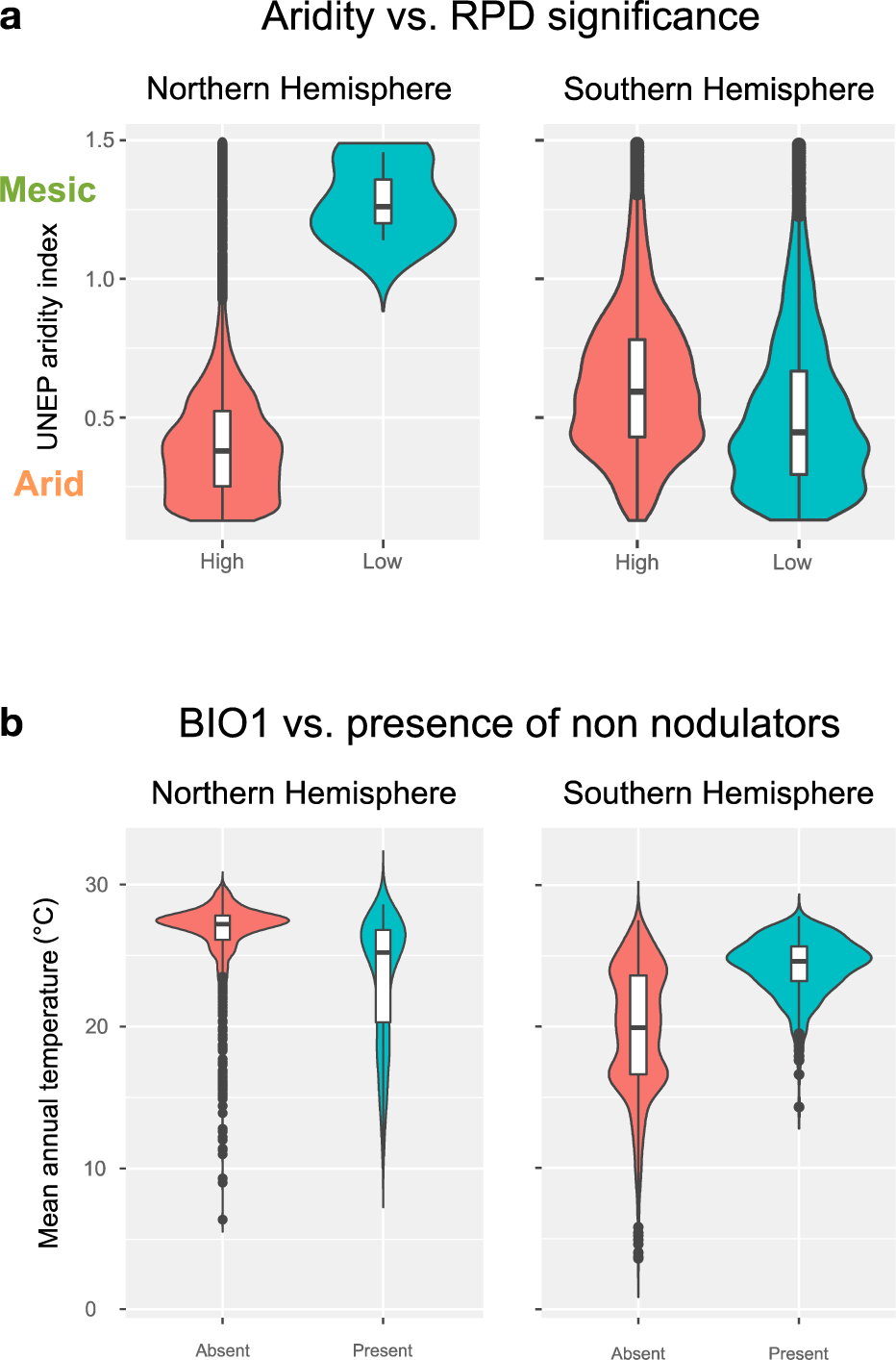
Violin plots of significance categories and non-nodulator presence, here shown as Northern-Southern Hemisphere contrasts. (A) Aridity vs. RPD significance, where left shows the Northern and right shows the Southern Hemisphere. (B) Mean annual temperature vs. non-nodulator presence, where left shows the Northern and right shows the Southern Hemisphere.

**Fig. 7:**
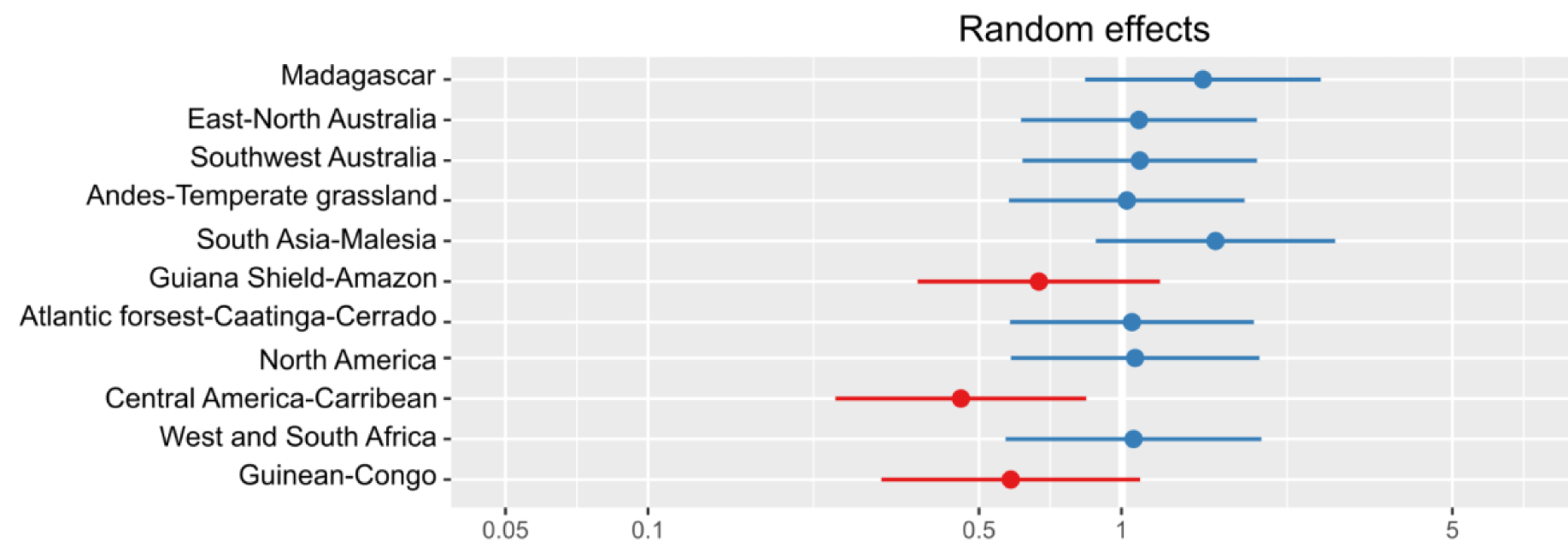
Random effect plot, showing the normalized effects of phyloregions on nodulating proportion, independent of environmental effects. Red effects indicate a negative effect (i.e., a substantial phyloregion effect increasing non-RNS species independent of environment).

## DISCUSSION

### Drivers of non-nodulator distributions

—This study represents to our knowledge the first quantitative investigation of the distribution of non-nodulating species in the largely nodulating legumes. The mimosoids form a prime clade to investigate the “tropical paradox” of significant biome and biogeographic regionalization of non-RNS strategies in largely RNS-associated clades (Sprent et al., 2017). We expected the RNS distribution to be subject to niche filtering, which would lead to its association with environmental factors thought to favor RNS. The distribution of non-RNS mimosoid lineages corresponded partly to expectations based on ecology (Sprent, 2007, 2009; Sprent et al., 2017; but see below): primarily moist tropical understory environments, especially in Africa. Accordingly, presence of non-RNS species was linked to habitats with higher overall precipitation, lower aridity, and lower temperature seasonality. However, Sprent’s mechanistic hypothesis that this is mediated by lack of N-limitation was not supported because soil N is a relatively unimportant predictor of the distribution of non-RNS species in our models. Mean annual temperature was not included in the best model for nodulation proportion, but this may be due to differing temperature responses in different biogeographic domains, as non-nodulators in the Northern Hemisphere were associated with cooler climates, but with warmer climates in the Southern Hemisphere (Fig. 6).

Overall, our results partially validated the observations of Sprent et al. (2017) that non-nodulating mimosoids are specifically associated with relatively moist habitats, even though some of these non-RNS species occur in drier habitats (e.g., the occurrence of non-nodulators in Central and South America). Griesmann et al. (2018) speculated that high nitrogen availability could promote the loss of RNS. Similarly, Sprent et al. (2017), although not explicitly differentiating loss vs. ancestral lack of non-RNS, clearly argued that moist environments that are not nitrogen-limiting are associated with absence of RNS. The model presented here is only partly congruent with the scenario that nitrogen availability drives non-RNS distributions; while high-nitrogen sites have more non-nodulating species, soil nitrogen content was not in the favored predictor set (Table S3), and therefore non-RNS distributions are better predicted by climate. As a corollary, it has recently been shown that tropical understory environments that are limited by light disfavor RNS and that this effect is stronger than soil nitrogen inputs (McCulloch & Porder, 2021; Taylor & Menge, 2018). This represents the first explicit test of the hypothesis that absence of RNS is specifically associated with moist tropical environments; future work should address this finding through replication in other nodulating clades and, as here we focused on a clade comprising only putative losses (Kates et al., 2022), particularly investigation of lineages possessing putative gains of nodulation.

### RNS and hotspots of mimosoid diversity

—Regions rich in non-RNS species were associated overall with significantly high RPD compared to null expectations; a strong RNS-RPD relationship has been previously observed in Fagales (Folk et al., 2023). Here we document a relationship between high RPD and non-RNS, whereas Folk et al. (2023) found a relationship between high RPD and RNS in Fagales (non-legume actinorhizal RNS species with a different bacterial symbiont). Much as Sprent (2017) predicted, equatorial Africa is the richest area in terms of non-RNS mimosoid species, and also has the greatest mimosoid phylogenetic diversity of anywhere on Earth. Africa also displays the strongest contrast between SR and RPD of any area for the mimosoids. Mexico and parts of eastern South America form the remainder of regions where significantly high RPD and high non-RNS mimosoid diversity align; only in the Amazon are non-RNS species found without RPD significance. Mexico, however, was the only area rich in non-RNS species that also shows substantial endemism. The other major RPD hotspots were Southeast Asia (including Malesia and a small southern area in South Asia) and eastern South America (an area comprising the Atlantic Forest, the Caatinga, and Cerrado in Brazil).

The association between relative phylogenetic diversity and RNS may be due to niche filtering (Cavender-Bares et al., 2004), in this instance representing environmental selection against RNS leading to an accumulation of species lacking RNS in areas that do not favor RNS strategies. Pointedly, in each of the areas where non-RNS species are found, co-occurring lineages that bear the non-RNS trait are distantly related and likely represent separate loss events (Kates et al., 2022). Africa has the greatest non-RNS diversity at the genus level (West Africa: *Aubrevillea, Calpocalyx, Tetrapleura*; only the small genus *Newtonia* inhabits both East and West Africa). While all early-diverging mimosoids, none of these co-occurring lineages are closely related. In much of the interior of Mexico and the northeast of South America, two distantly related non-RNS genera occur: *Zapoteca*, endemic to the Americas, and *Parkia*, a pantropical genus. Thus, our results indicate that the convergent loss of RNS in mimosoids, represented by at least four separate loss events (Kates et al., 2022), is associated with tropical relatively moist forest understories. In every instance, non-RNS communities harbor high mimosoid phylogenetic diversity and generally include multiple distantly related non-RNS genera. While multiple explanations can be invoked for the isolated observation of high RPD, alternative hypotheses, such as a local radiation of non-RNS species, do not explain sympatry of distinct lineages bearing the same trait, underlining environmental filtering against RNS as the best explanation given the present data.

Consistent with the contrasting non-RNS lineages in each non-RNS hotspot, lack of RNS was found in five distinct phyloregions: the three in the Americas were the Mexico-Caribbean region, the Guiana Shield-Amazon, and the Atlantic Forest-Caatinga-Cerrado. In Africa, both phyloregions (Guinean-Congo and west and south African tropical dry forest) contained non-RNS species. Based on the phylogenetic regionalization dendrogram, these five regions belong to two separate continent-level phylogenetic clusters. While non-RNS species are most numerous and spatially extensive in rainforest, more puzzling is the occurrence of non-RNS species in both the African tropical dry forest, Mexico, and the northeast of South America, areas that do not perfectly represent the moist tropical forest model of Sprent et al. (2017) and are an exception to the patterns highlighted here. The three involved genera, *Zapoteca, Parkia,* and *Newtonia*, are habitat generalists that occupy both moist and dry tropical forest. No mimosoid lacking RNS is completely specialized to dry tropical environments because all lineages are polymorphic with respect to precipitation niche; thus, the outlier areas could therefore represent habitat shifts after loss of RNS. Therefore, while we found strong evidence for present-day associations between precipitation and RNS status, it remains to be investigated whether the non-RNS genera originated in similar ecological contexts.

The only major areas with significantly low RPD, none aligned with non-RNS distributions, were the Southwest and especially East Australia hotspots, consistent with the idea that Australian mimosoid diversity is the result of recent rapid diversification (Dale et al., 2020; Miller et al., 2013; Renner et al., 2020). Australia is indeed the only extensive area on Earth with low mimosoid RPD when assessed at the global level. Endemism analyses demonstrated that the East Australia hotspot contained both neo- and mixed endemism, but the Southwest Australia hotspot was a center of mixed endemism, consistent with the general recognition of this flora as older than the flora on the rest of the Australian continent. These results are partly consistent with a focused phylogenetic diversity study in *Acacia* (Mishler et al., 2014) in recovering an age gradient across Australia, but Miller et al. (2013) found significant paleoendemism and high RPD in Southwest Australia, and mixed/neo-endemism in East Australia. It is important to consider that while the taxonomic focus in Australia is largely the same between the two studies because *Acacia* dominates the mimosoid flora of Australia, the scope of comparison is different in our global study. Similarly, spatial scale may be responsible for the contrast between our results and the main finding of Dale et al. (2020) that species distributions do not follow recognized biomes; our regionalization map closely resembles their biome map (Dale et al., 2020: Fig. 1), and centers of endemism were identified in all three major regions, which may reflect the global analysis scale. Three further very small areas of significantly low RPD and two areas of paleoendemism were identified in South America; these are likely artifactual as they represent transition areas between major biomes and could have been driven by overlapping edge ranges of endemic species.

### Conclusions

—Mimosoids are distributed in nine distinct diversity hotspots on the basis of SR, which overlap with major recognized tropical dry forest hotspots. Only about half of these, including both moist and seasonally dry forest, are hotspots of high RPD; only two are clear hotspots of low RPD and represent recent rapid radiations of the large Australian mimosoid flora. Lineages lacking RNS display two major hotspots in Africa and the Americas, which phylogenetic regionalizations identified as two biogeographically independent domains. The proportion of non-RNS species is strongly linked to high RPD, suggesting the operation of niche filtering. As an additional line of evidence for niche filtering in lowland tropical forest, close examination reveals that the richest non-RNS areas are characterized by sympatry of highly disparate non-RNS lineages, different in each phyloregion, that represent independent evolutionary origins of non-RNS. Our work validates the hypothesis that non-RNS mimosoids are specifically associated with tropical environments that are relatively moist and high in nitrogen, areas where nitrogen availability and investment may not favor RNS strategies.

## ACKNOWLEDGMENTS

This work was funded by the National Science Foundation, DEB-1916632.

## DATA ACCESSIBILITY STATEMENT

Scripts to perform the modeling analyses and raw grid cell products have been deposited in GitHub (https://github.com/ryanafolk/mimosoid_phylodiversity).

