## Supplement for "Phylogenetic diversity and regionalization of root nodule symbiosis"

**Supplemental table S1.** Summary of percent of species sampled per genus. Denominators assume WCVP accepted species.

| Genus (following LPWG) | Percent sampled: phylogeny | Percent sampled: SDM |
|---|---:|---:|
| ***Abarema*** | 76.1% | 63.0% |
| ***Acacia*** | 57.3% | 49.8% |
| ***Acaciella*** | 33.3% | 13.3% |
| ***Adenanthera*** | 33.3% | 33.3% |
| ***Adenopodia*** | 42.9% | 42.9% |
| ***Afrocalliandra*** | 0.0% | 0.0% |
| ***Alantsilodendron*** | 11.1% | 11.1% |
| ***Albizia*** | 53.7% | 44.7% |
| ***Amblygonocarpus*** | 0.0% | 0.0% |
| ***Anadenanthera*** | 50.0% | 50.0% |
| ***Archidendron*** | 17.3% | 14.3% |
| ***Archidendropsis*** | 21.4% | 21.4% |
| ***Aubrevillea*** | 50.0% | 50.0% |
| ***Balizia*** | 66.7% | 66.7% |
| ***Blanchetiodendron*** | 100.0% | 100.0% |
| ***Calliandra*** | 44.0% | 29.3% |
| ***Calliandropsis*** | 100.0% | 0.0% |
| ***Calpocalyx*** | 36.4% | 36.4% |
| ***Cathormion*** | 100.0% | 200.0% |
| ***Cedrelinga*** | 0.0% | 100.0% |
| ***Chidlowia*** | 100.0% | 100.0% |
| ***Chloroleucon*** | 27.3% | 0.0% |
| ***Cojoba*** | 66.7% | 46.7% |
| ***Cylicodiscus*** | 100.0% | 100.0% |
| ***Desmanthus*** | 45.8% | 41.7% |
| ***Dichrostachys*** | 12.5% | 12.5% |
| ***Ebenopsis*** | 66.7% | 0.0% |
| ***Elephantorrhiza*** | 37.5% | 37.5% |
| ***Entada*** | 31.3% | 18.8% |
| ***Enterolobium*** | 63.6% | 36.4% |
| ***Faidherbia*** | 100.0% | 100.0% |
| ***Falcataria*** | 33.3% | 33.3% |
| ***Fillaeopsis*** | 0.0% | 100.0% |

| | | |
|---|---|---|
| ***Gagnebina*** | 60.0% | 60.0% |
| ***Havardia*** | 57.1% | 0.0% |
| ***Hesperalbizia*** | 100.0% | 100.0% |
| ***Hydrochorea*** | 50.0% | 50.0% |
| ***Indopiptadenia*** | 0.0% | 0.0% |
| ***Inga*** | 53.1% | 42.2% |
| ***Jacqueshuberia*** | 57.1% | 42.9% |
| ***Kanaloa*** | 100.0% | 0.0% |
| ***Lemurodendron*** | 100.0% | 100.0% |
| ***Leucaena*** | 40.0% | 16.0% |
| ***Leucochloron*** | 0.0% | 0.0% |
| ***Lysiloma*** | 87.5% | 25.0% |
| ***Macrosamanea*** | 36.4% | 36.4% |
| ***Mariosousa*** | 0.0% | 0.0% |
| ***Microlobius*** | 100.0% | 0.0% |
| ***Mimosa*** | 30.9% | 19.8% |
| ***Mimozyganthus*** | 0.0% | 0.0% |
| ***Neptunia*** | 50.0% | 25.0% |
| ***Newtonia*** | 25.0% | 18.8% |
| ***Painteria*** | 0.0% | 0.0% |
| ***Parapiptadenia*** | 33.3% | 16.7% |
| ***Pararchidendron*** | 20.0% | 20.0% |
| ***Paraserianthes*** | 100.0% | 100.0% |
| ***Parkia*** | 25.0% | 7.5% |
| ***Pentaclethra*** | 66.7% | 33.3% |
| ***Piptadenia*** | 39.3% | 14.3% |
| ***Piptadeniastrum*** | 100.0% | 100.0% |
| ***Piptadeniopsis*** | 100.0% | 100.0% |
| ***Pithecellobium*** | 87.5% | 58.3% |
| ***Pityrocarpa*** | 66.7% | 66.7% |
| ***Plathymenia*** | 100.0% | 100.0% |
| ***Prosopidastrum*** | 28.6% | 14.3% |
| ***Prosopis*** | 49.1% | 28.1% |
| ***Pseudopiptadenia*** | 18.2% | 9.1% |
| ***Pseudoprosopis*** | 42.9% | 42.9% |
| ***Pseudosamanea*** | 0.0% | 0.0% |
| ***Samanea*** | 40.0% | 40.0% |
| ***Sanjappa*** | 0.0% | 0.0% |
| ***Schleinitzia*** | 0.0% | 25.0% |

| | | |
|---|---|---|
| ***Senegalia*** | 5.3% | 10.6% |
| ***Serianthes*** | 11.8% | 5.9% |
| ***Sphinga*** | 100.0% | 200.0% |
| ***Stryphnodendron*** | 28.0% | 12.0% |
| ***Tetrapleura*** | 100.0% | 100.0% |
| ***Tetrapterocarpon*** | 100.0% | 100.0% |
| ***Thailentadopsis*** | 0.0% | 0.0% |
| ***Vachellia*** | 3.1% | 4.3% |
| ***Viguieranthus*** | 27.8% | 27.8% |
| ***Wallaceodendron*** | 100.0% | 100.0% |
| ***Xerocladia*** | 100.0% | 100.0% |
| ***Xylia*** | 33.3% | 33.3% |
| ***Zapoteca*** | 30.4% | 21.7% |
| ***Zygia*** | 31.7% | 6.7% |

**Table S2.** Model choice. Best AIC in boldface.

| Model | RPD AIC | CANAPE AIC | Proportion nodulator AIC |
|---|---|---|---|
| **GLM, full model** | 302077.9 | 14686.77 | -494090.8 |
| **GLM, simple model** | 303263.8 | 14794.27 | -499397.2 |
| **LMM, full model** | **235183.4** | **5210.77** | **-498329.2** |
| **LMM, simple model** | 239056.7 | 5249.756 | -503749.9 |
| **LMM, no-environment model** | 269709.9 | 6224.892 | -477423 |
| **ΔAIC** | 3873.3 | 38.986 | 5420.7 |

**Table S3.** Model parameters. Highest normalized coefficient in boldface.

| Model predictor | Best RPD model normalized coefficient | Best CANAPE model normalized coefficient | Best proportion nodulation normalized coefficient |
|---|---|---|---|
| **UNEP aridity index** | -0.119314 | -1.67218 | 0.155088 |
| **Bioclim 1** | -0.016681 | 2.29839 | n/a |
| **Bioclim 12** | 0.037962 | -0.75828 | **-0.224597** |
| **Bioclim 7** | **-0.265119** | 1.08990 | 0.101997 |
| **Bioclim 17** | 0.040996 | 0.41246 | n/a |
| **Nitrogen content** | 0.005390 | -0.47975 | n/a |
| **pH** | -0.260911 | **-2.62390** | 0.168325 |
| **Organic carbon content** | 0.063158 | -0.50673 | 0.077060 |
